# TOPOVIBL function in meiotic DSB formation: new insights from its biochemical and structural characterization

**DOI:** 10.1101/2023.11.02.565342

**Authors:** Boubou Diagouraga, Izabella Tambones, Coralie Carivenc, Chérine Bechara, Bernard de Massy, Albane le Maire, Thomas Robert

## Abstract

The TOPOVIL complex catalyzes the formation of DNA double strand breaks (DSB) that initiate meiotic homologous recombination, an essential step for chromosome segregation and genetic diversity during gamete production. TOPOVIL is composed of two subunits (SPO11 and TOPOVIBL) and is evolutionarily related to the archaeal TopoVI topoisomerase complex. SPO11 is the TopoVIA subunit orthologue and carries the DSB formation catalytic activity. TOPOVIBL shares homology with the TopoVIB ATPase subunit. TOPOVIBL is essential for meiotic DSB formation, but its molecular function remains elusive, partly due to the lack of biochemical studies. Here, we purified TOPOVIBLΔC25 and characterized its structure and mode of action *in vitro*. Our structural analysis revealed that TOPOVIBLΔC25 adopts a dynamic conformation in solution and our biochemical study showed that the protein remains monomeric upon incubation with ATP, which correlates with the absence of ATP binding. Moreover, TOPOVIBLΔC25 interacted with DNA, with a preference for some geometries, suggesting that TOPOVIBL senses specific DNA architectures. Altogether, our study identified specific TOPOVIBL features that might help to explain how TOPOVIL function evolved toward a DSB formation activity in meiosis.

## Introduction

In most organisms, the accurate segregation of the homologous chromosomes during the first meiotic division depends on homologous recombination. This is initiated by the programmed formation of DNA double strand breaks (DSBs) that is catalyzed by the conserved TOPOVIL complex and by its accessory factors (1–3). Eukaryotic TOPOVIL is composed of two subunits that are both essential for meiotic DSB formation: SPO11 and TOPOVIBL (4, 5) (3, 6). The meiotic TOPOVIL complex is evolutionarily linked to the archaeal TopoVI type IIB topoisomerase (7) that regulates DNA topology through the formation of transient DNA DSBs (8). The archaeal TopoVI is composed of two subunits: TopoVIA and B (the orthologues of SPO11 and TOPOVIBL, respectively) that assemble as an A_2_/B_2_ hetero-tetramer (9, 10). The A subunit is the catalytic part of the enzyme and the B subunit corresponds to the ATP binding region. Structural studies suggest that TopoVI relaxation activity is mainly driven by the B subunit motion, triggered by ATP binding/hydrolysis and interaction with DNA. It was proposed that the reaction cycle initiates by the binding of one DNA duplex (G DNA segment) to the A subunit dimer (11). This is followed by the capture of a second duplex (T DNA segment) upon ATP binding by the B subunit that triggers its dimerization and the G segment cleavage to form a DSB with a covalent protein-DNA link (12, 13). Then, ATP hydrolysis induces the B subunit motion that triggers the separation of the A dimer, the opening of the G segment and the passage of the T segment through the break. This promotes G segment religation followed by the enzyme reset through ADP release (9, 14). Three regions of the B subunit interact with DNA: the KGRR loop, the Stalk/WKxY motif, also named WKxY-containing octapeptide (WOC) motif, and the H2TH domain. The interaction with the first two motifs is essential for TopoVI function. Indeed, it was proposed that the KGRR loop helps DNA crossing recognition and ATP turnover, while the WOC motif is important for G segment binding (15).

In meiosis, TOPOVIL achieves a partial TopoVI cycle and its activity is frozen at the DSB formation step. *In vivo* data showed that DSBs are generated through the formation of concerted double nicks catalyzed by SPO11, which remains covalently bound to the DNA ends (3, 6)(4, 5, 16). This mechanism leads to a non-catalytic reaction, with the absence of the religation step and a suicide reaction for TOPOVIL. This meiotic activity implies similarities and differences between TOPOVIL and the TopoVI topoisomerase. Particularly, like the TopoVIA subunit, SPO11 is believed to act as a dimer, suggesting the formation of an active hetero-tetramer composed of two SPO11 and two TOPOVIBL (17–19)(1, 2). TOPOVIBL displays only partial homology with TopoVIB. Indeed, TopoVIB contains a GHKL/Bergerat fold, a H2TH motif, the transducer domain and a disordered C-terminal domain (CTD) (9, 10). Conversely, TOPOVIBL harbors a degenerate GHKL/Bergerat fold-ATPase domain, a linker domain replaces the H2TH DNA interacting domain, the central transducer domain is present but, the disordered CTD is divergent (Fig. 1*A*) (3, 6). TOPOVIBL molecular function remains puzzling, particularly its potential ATP-dependent dimerization and interaction with DNA, which are pivotal for the TopoVI cycle.

**Fig. 1.**
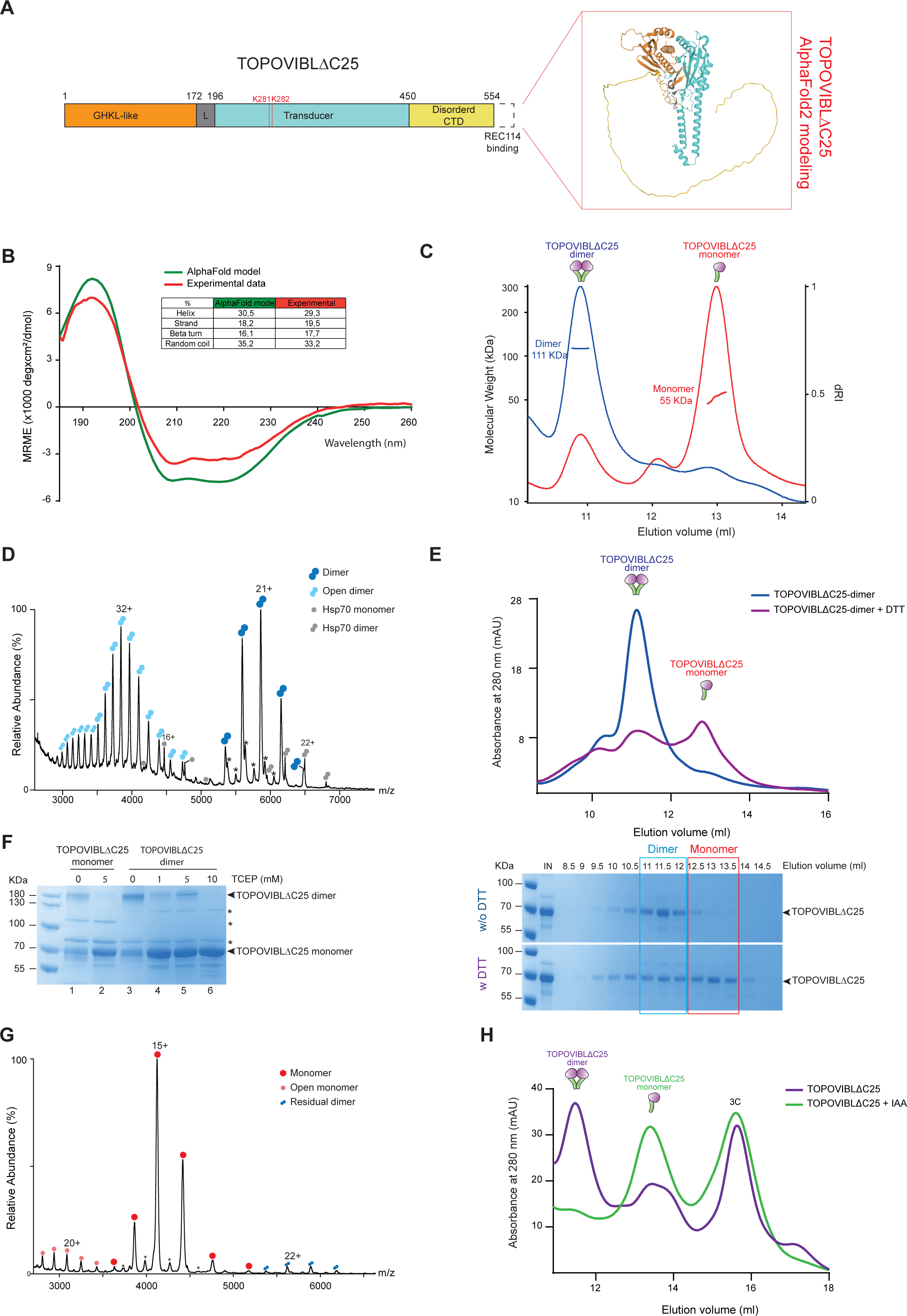
TOPOVIBLΔC25 purification and characterization. **(*A*)** Schematic representation of TOPOVIBLΔC25. Red box: 3D structure model of TOPOVIBLΔC25 by the AlphaFold2 Protein Structure Database. **(*B*)** Circular dichroism analysis of TOPOVIBLΔC25 secondary structure. **(*C)*** SEC-MALS analysis of TOPOVIBLΔC25 monomers (red) and dimers (blue). **(*D*)** Native MS spectrum of TOPOVIBLΔC25 homodimer peaks at 1.5 mg/ml showing the presence of two charge state distributions of the dimeric species: one at an average charge state of 21+ corresponding to an average MW of 123,044 ± 24 Da (double dark blue circles), and another at an average charge state of 32+ corresponding to an average MW of 122,908 ± 76 Da (double light blue circles). As the dimer theoretical MW is 122,639 Da, the observed additional mass relates to residual adducts that remain bound to the protein in native conditions. Asterisks indicate hydrolyzed TOPOVIBLΔC25 at 118,000 and 121,000 Da. Some contaminant species are also present at low intensity, identified by proteomics as monomeric (71,448 ± 7 Da, gray circles) and homodimeric HSP70 (142,875 ± 7 Da, double gray circles), see Table S1. **(*E*)** SEC analysis of TOPOVIBLΔC25 dimer incubated with DTT. Upper panel: chromatograms of the elution profiles of the dimer (blue) and of the DTT-treated dimer (purple). Lower panel: SDS-PAGE of the SEC elution fractions (8.5 ml to 14.5 ml) of TOPOVIBLΔC25 dimers not incubated (top gel) or incubated with DTT (bottom gel). IN: SEC input. **(*F*)** Non-reducing-PAGE analysis followed by Coomassie staining of TOPOVIBLΔC25 dimers and monomers incubated with TCEP (a reducing agent). *: contaminants. **(*G*)** Native MS spectrum of blocked TOPOVIBLΔC25 monomers (1.5 mg/ml) showing the presence of a main distribution with an average charge state of 15+ corresponding to an average MW of 61,844 ± 55 Da (red circles) and another distribution corresponding to a highly charged monomer with an average MW of 61,710 ± 8 Da (light red circles). The theoretical MW of the monomer with all cysteines reduced and no carbamidomethylation is 61,319 Da. Some residual TOPOVIBLΔC25 homodimers are also present at low intensity. Asterisks indicate hydrolyzed TOPOVIBLΔC25 at 59 kDa. **(*H*)** Superimposition of the SEC elution profile chromatograms of TOPOVIBLΔC25 purified with (green curve) or without 10 mM IAA in the cell lysis buffer (purple curve). The peaks corresponding to TOPOVIBLΔC25 dimers and monomers are indicated.

In *Saccharomyces cerevisiae*, the identified TOPOVIBL orthologue Rec102 shares similarity only with the transducer domain. It was proposed that the Rec104 protein replaces the GHKL domain and that the Rec102/Rec104 complex fulfills TOPOVIBL function (6, 7, 20–22). The yeast meiotic DSB core complex was defined *in vitro* as a 1:1:1:1 stoichiometry complex, composed of Spo11, Rec102, Rec104 and Ski8 (20–22). This core complex interacts with DNA, and Rec102 is one of the contact points. However, the core complex does not dimerize *in vitro* and consequently is not active for DSB formation. Therefore, the molecular function of Rec102/Rec104 (TOPOVIBL counterpart) in these processes remains to be understood. Recently, it was proposed that the yeast Rec114, Mei4 and Mer2 DSB accessory factors assemble in a complex. This complex forms condensate-like structures in the presence of DNA that directly interacts with Rec102/Rec104 and thus might favor the core complex dimerization (23–26)(27).

The different domains composing TopoVIB have been identified in the mouse TOPOVIBL sequence. However, based on AlphaFold2 modeling of TOPOVIBL (AF-J3QMY9-F1-model), we recently showed that the ATP binding site of the GHKL domain is degenerated and lacks the ATP-lid structure and the N-terminal strap region involved in the ATP-dependent dimerization, suggesting a meiotic-specific regulation of the TOPOVIBL subunit (6, 6, 7). We also found that TOPOVIBL directly interacts with the mouse REC114 DSB accessory protein and that this interaction is essential for DSB formation in some genomic contexts (28). These results indicate that TOPOVIBL regulates, in part, the TOPOVIL DSB catalytic activity through protein-protein interactions. However, how TOPOVIBL is intrinsically regulated at the molecular level to promote the DSB formation activity of the TOPOVIL complex rather than the cutting/resealing activity of its archaeal counterpart remains unknown.

A biochemical characterization of the meiotic B subunit to understand the regulation of this essential activity for sexual reproduction is missing, probably due to difficulties in purifying the soluble protein. Here, we present a biochemical and structural characterization of the mouse TOPOVIBL subunit, which represents a first important step in the characterization of TOPOVIBL role in TOPOVIBL-SPO11 mode of action. We purified a soluble and structured TOPOVIBL form that lacks the last 25 C-terminal residues: TOPOVIBLΔC25. We showed that this protein adopts a dynamic conformation in solution, in equilibrium between an extended and a compact form. Unlike its archaeal counterpart, it did not dimerize upon incubation with ATP and DNA, and did not interact with ATP. However, we found that TOPOVIBLΔC25 interacts with DNA, with a topological preference for linear single-stranded (ss) DNA and negative supercoiled DNA. This highlights the importance of TOPOVIBL interaction with some DNA geometries for TOPOVIL regulation. In addition, the TOPOVIBLΔC25-DNA interaction only partly relied on the known conserved WOC motif, suggesting the presence of additional contact points with DNA. From this study, we identified common and divergent features with the archaeal TopoVIB that might help to explain how TOPOVIL function evolved toward a DSB formation activity in meiosis.

## Results

### Purification of mouse TOPOVIBLΔC25

To characterize the molecular mechanisms underlying TOPOVIBL activity, we purified *in vitro* the mouse TOPOVIBL protein. As full length TOPOVIBL fused to the maltose binding protein (MBP) tag was mostly insoluble after expression in Sf9 insect cells, we tested different truncated versions. TOPOVIBLΔC25, which lacks the last 25 C-terminal residues that fold into a single helix essential for the interaction with the REC114 meiotic protein (Fig. 1*A*) (28), showed the highest solubility. AlphaFold2 structure prediction (AF-J3QMY9-F1-model) suggested that TOPOVIBLΔC25 includes a globular domain at its N-terminus that contains the GHKL-Like and Linker motifs (respectively residues 1 to 172 and 172 to 196), followed by the transducer domain (a long alpha-helix, residues 196 to 450) that is connected to a disordered region of 119 amino acids (Fig. 1*A*). Among the different tested constructs, TOPOVIBLΔC25 was the closest in structure to the full-length protein. Moreover, we previously showed by yeast two-hybrid assay that it interacts with SPO11 (28), suggesting that it is correctly folded *in vivo*. Therefore, we decided to characterize in vitro TOPOVIBLΔC25 that we consider the most informative to assess the TOPOVIBL subunit role.

We affinity-purified TOPOVIBLΔC25 fused to the MBP tag, and then released the protein from the resin by tag cleavage with the PreScission protease followed by purification by size exclusion chromatography (SEC) (Fig. S1*A-B*). We confirmed by tandem mass spectrometry (MS/MS) that the purified protein corresponded to TOPOVIBLΔC25: 40 unique peptides identified all along the protein and the highest iBAQ value (Fig. S1*C* and Table S1). Far-UV circular dichroism analysis of purified TOPOVIBLΔC25 confirmed the presence of secondary structures in a proportion that matched the AlphaFold2 model derived from AF-J3QMY9-F1 pdb file (Fig. *1B*), indicating proper folding of purified TOPOVIBLΔC25.

During SEC purification of TOPOVIBLΔC25, we observed a second peak at a lower elution volume that could correspond to the protein eluted as homodimer (Fig. S1*B*). To better characterize the different purified species, we used SEC coupled to multiangle light scattering (SEC-MALS). The experimental molecular weight (MW) values were compatible with two oligomeric states: 55.1 ± 0.15 kDa for the monomer and 111.1 ± 0.007 kDa for the homodimer (Fig. 1*C*), compared with the theoretical MW of 61.32 kDa and 122.64 kDa for the monomeric and dimeric form.

We further characterized the protein in its putative homodimer state by native MS that gave a MW of 123.044 ± 0.024 KDa, in accordance with the expected MW of a homodimer (Fig. 1*D*). In native MS spectra, we observed systematically two charge state distributions that corresponded to the homodimer mass, one with an average charge of 21+ and another with a higher average charge of 32+ (Fig. 1*D*). This suggests that TOPOVIBLΔC25 exists in a closed and a more open conformation. The second could be related to the disordered region in the protein and/or to unfolded TOPOVIBLΔC25 species. Increasing the activation during MS acquisition did not lead to the disruption of the dimeric stoichiometry. This implied the presence of a covalent interaction between monomers. As TOPOVIBLΔC25 contains eleven cysteines, we hypothesized that these residues can form disulfide bonds during protein production or purification. To test this hypothesis, we incubated TOPOVIBLΔC25 homodimers with 50 mM dithiothreitol (DTT, a reducing agent), followed by SEC analysis. We observed a displacement of the homodimer toward the monomeric form (Fig. 1*E*). Similarly, incubation of the homodimer with different concentrations of tris(2-carboxyethyl)phosphine (TCEP), from 1 mM to 10 mM, followed by protein separation on non-reducing acrylamide gels showed that homodimers can be fully converted into monomers (Fig. 1*F*). Moreover, native MS analysis of SEC-purified TOPOVIBLΔC25 homodimers that were subsequently incubated with 50 mM DTT and 10 mM 2-Iodoacetamide (IAA), which alkylates the accessible cysteine residues and prevents the formation of disulfide bonds, confirmed the formation of stable blocked monomers with a measured MW of 61,844 ± 55 Da (Fig. 1*G* and Table S2). These results show that TOPOVIBLΔC25 homodimers result from disulfide bond formation.

Lastly, to evaluate whether these bonds are induced by the oxidative environment of the cell lysis conditions or are formed during protein folding in Sf9 cells, we purified TOPOVIBLΔC25 in lysis buffer that included 10 mM IAA. IAA alkylated and blocked the free cysteines, but not the cysteines that were engaged in disulfide bonds in the cells *in vivo* during protein expression and folding. SEC showed that in this condition, TOPOVIBLΔC25 was mainly purified as monomers (Fig. 1*H* and Fig. S1*D*). Therefore, we propose that i) TOPOVIBLΔC25 homodimer formation is due to stochastic intermolecular interactions between cysteines, favored by the oxidative environment of the purification conditions; and ii) TOPOVIBLΔC25 homodimers have no physiological relevance. In line with these assumptions, AlphaFold2 did not generate homodimers. Therefore, we focused on the monomeric form.

### TOPOVIBLΔC25 monomers adopt a dynamic conformation in solution

To evaluate the conformation of TOPOVIBLΔC25 monomers in solution, we performed SEC coupled to Small-Angle X-ray Scattering (SEC-SAXS) that provides low resolution structural information (Fig. 2*A*). SEC-SAXS data indicated that TOPOVIBLΔC25 had a radius of gyration (Rg) of 3.65 ± 0.06 nm and a maximum intramolecular distance (Dmax) of 15.6 ± 0.6 nm in solution. MW estimation confirmed that TOPOVIBLΔC25 remained as monomers (experimental MW of 62.35 kDa vs theoretical MW of 61.32 kDa) in the SEC-SAXS experimental conditions. The dimensionless Kratky plot does not display a perfect bell-shaped peak but rather a combination of a bell-shape and a plateau at higher gRg values and displays a shift from the values expected for globular proteins on both *x* and *y* axes (29), indicative of flexibility in the protein. The asymmetrical pairwise distance distribution function, P(r), confirmed TOPOVIBLΔC25 flexibility (Fig. 2*B*, inset). To precisely characterize the protein structure, we generated an ensemble of conformations based on the AlphaFold2 model of TOPOVIBLΔC25, especially for the protein core that comprises the GHKL-like domain, the linker and the transducer motif. The C-terminal domain (from C451 to S579) was built as a random-coil region. Fitting the SAXS profile using the Ensemble Optimization Method (EOM) (30) indicated that TOPOVIBLΔC25 C-terminal domain did not behave as a fully random coil part. Indeed, the SAXS profile of TOPOVIBLΔC25 was best described with an excellent fit to the experimental profile (chi-square of 0.76) by EOM selected sub-ensembles showing that the C-terminal region adopts two distinct conformations leading to an elongated form of TOPOVIBLΔC25 (with Rg>40Å), where the C-terminal region is highly extended and to a more compact form of TOPOVIBLΔC25 (with Rg<35Å), where the C-terminal region is in close proximity to the core of the protein. The latter represents 66% of the selected conformations over 20% frequency of extended (Fig. 2*C*). Importantly, the peaks corresponding to these two forms were robust relative to the parameters used for the genetic algorithm optimization in EOM and supported the hypothesis that TOPOVIBLΔC25 exists in a dynamic conformational equilibrium between two main states, with a preference for the compact form in solution (Fig. 2*D*). This suggests an allosteric regulation of the TOPOVIBL subunit.

**Fig. 2.**
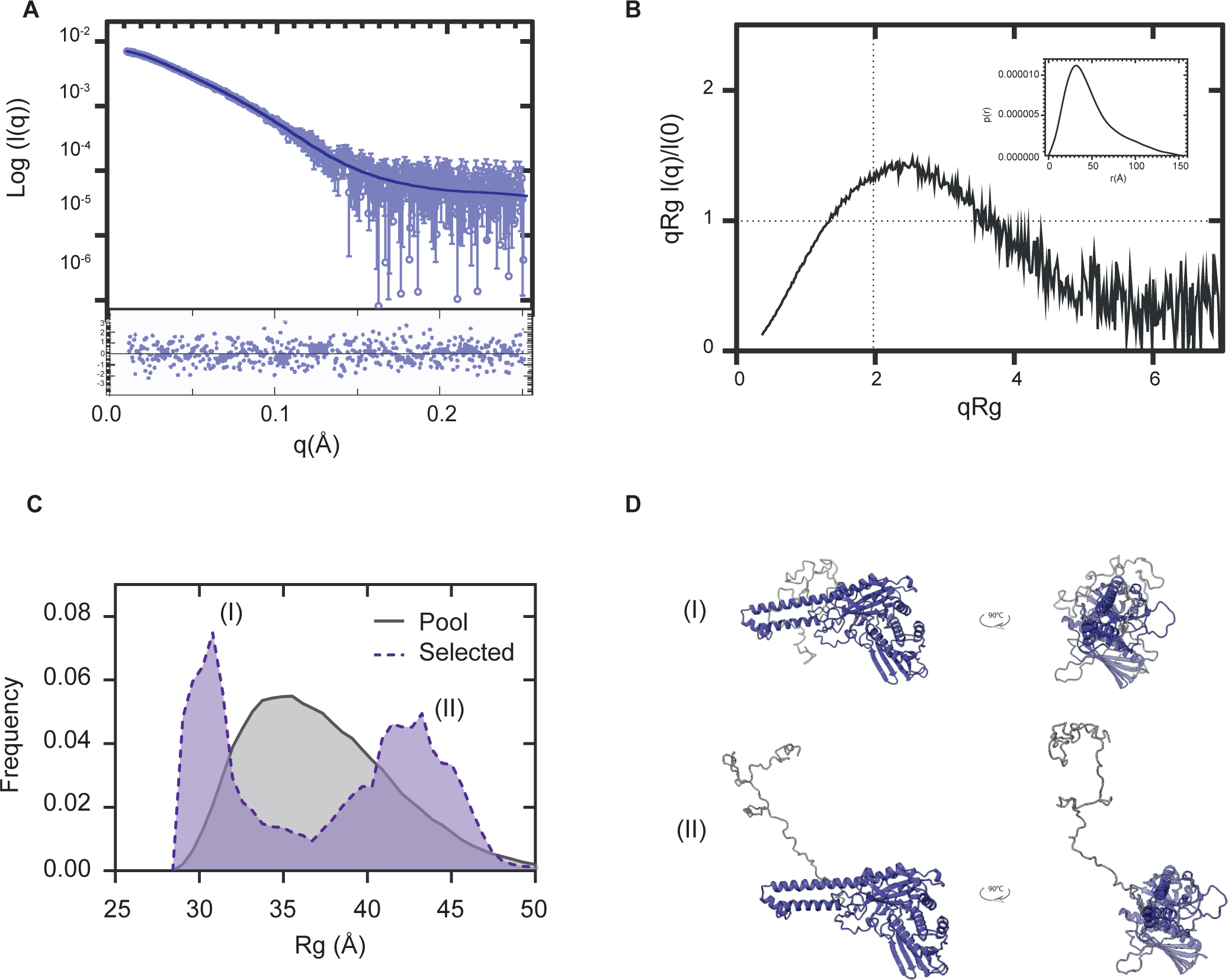
SAXS data shows a flexible, but preferential conformation states of TOPOVIBLΔC25 in solution. **(A)** The SAXS intensity profile for monomeric TOPOVIBLΔC25 (open circles) is overlaid with the theoretical curve derived from the EOM sub-ensemble selection (solid line). Residuals from the EOM fitting are shown below. **(B)** The dimensionless Kratky plot calculated from the SAXS profiles shows a shift from the values expected for globular proteins (shown as black dashed lines) on both the x and y axes. This and the asymmetrical form of the pair-distance distribution functions P(r) (inset) indicate the presence of flexibility in TOPOVIBLΔC25. **(C)** Radius of gyration (Rg) distribution of randomly generated ensembles (gray) and selected ensembles (purple), indicating a higher frequency (∼66%) of the compact conformation (i) than the elongated conformation (ii) of TOPOVIBLΔC25 in solution. **(D)** Representative TOPOVIBLΔC25 compact (i, Rg<35) and elongated states (ii, Rg>40) selected by EOM.

### TOPOVIBLΔC25 is not sensitive to ATP

In Archaea, ATP-induced TopoVIB dimerization is essential for TopoVI cutting and resealing activity (31). We previously reported that the ATPase domain of mouse TOPOVIBL is degenerated compared to its archaeal ancestor (28). Yet, despite the absence of the conserved N-strap dimerization domain in the TOPOVIBL sequence, we did not rule out the possibility of an ATP-dependent TOPOVIBLΔC25 dimerization. To address this question, first, we purified TOPOVIBLΔC25 in the presence of IAA to favor the formation of blocked monomers, followed by incubation, or not, with an excess of ATP. Both conditions led to the same SEC elution profile, compatible with the presence of stable monomers. This suggested that TOPOVIBL does not dimerize upon incubation with ATP (Fig. 3*A* and Fig. S2). To rule out the possibility of a rapid ATP turnover that might mask the ATP binding-dependent TOPOVIBL dimerization, we used an excess of AMP-PNP, the non-hydrolyzable ATP analogue. Again, we did not detect TOPOVIBL dimerization (Fig. 3*B*).

**Fig. 3.**
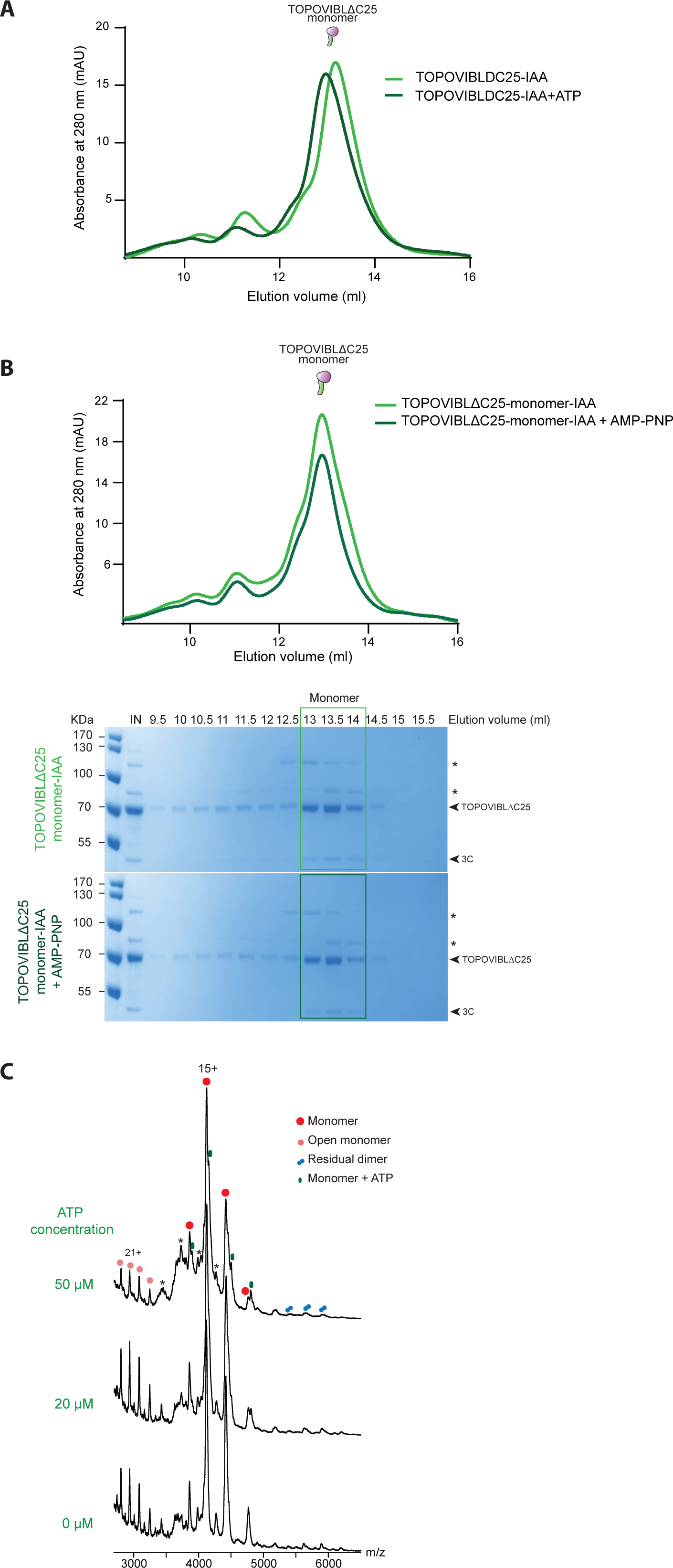
TOPOVIBLΔC25 is not sensitive to ATP and AMP-PNP. **(*A*)** SEC chromatograms of blocked TOPOVIBLΔC25 monomers incubated (dark green) or not (light green) with ATP. **(*B*)** SEC chromatograms of TOPOVIBLΔC25 monomers incubated or not with AMP-PNP. Upper panel: chromatograms of the elution profiles of TOPOVIBLΔC25 monomers not incubated (light green) and incubated with AMP-PNP (dark green). Lower panel: SDS-PAGE and Coomassie staining of the SEC elution fractions of TOPOVIBLΔC25 monomers not incubated (top gel) and incubated with AMP-PNP (bottom gel). The light green box and dark green box highlights the fractions corresponding to the monomer peak. IN: SEC input. *: contaminant bands, 3C: band of the PreScission protease. **(*C*)** Native MS analysis of 20 µM TOPOVIBLΔC25 monomers incubated with increasing amounts of ATP, showing the presence of rare, non-specific ATP binding at high concentrations (green circles). Red and light red circles represent blocked monomeric species, double blue circles represent residual dimeric species, and asterisks indicate contaminant species.

Second, we incubated blocked TOPOVIBLΔC25 monomers with two different ATP concentrations (20 µM and 50 µM, 1:1 and 1:2.5 molar ratio, respectively) before native MS analysis that did not highlight any strong interaction of TOPOVIBLΔC25 with ATP (Fig. 3*C*). The small ATP adducts observed did not increase significantly upon ATP addition, and were mainly visible at the lowest charge states of TOPOVIBLΔC25 (15+). This implies that the observed adducts are most probably related to non-specific binding occurring during the electrospray process. On the other hand, ATP addition did not induce any increase in the proportion of homodimeric species detected in the native MS spectra.

Our results suggest that TOPOVIBLΔC25 does not interact with ATP and experimentally validated our previous *in silico* predictions (28). Moreover, they show that unlike its archaeal orthologue, the TOPOVIBL subunit does not retain the ATP-dependent dimerization activity.

### TOPOVIBLΔC25 binds to DNA, with topological preferences

Archaeal TopoVI relaxation activity requires TopoVIB interaction with DNA, with a topological preference for linear duplex DNA of 60 bp, negative supercoils, and DNA crossings (15). To determine whether the meiotic TOPOVIBL subunit also interacted with DNA, independently of the presence of SPO11 (the TopoVIA orthologue), we first measured TOPOVIBLΔC25 affinity for DNA duplexes of different lengths and sequences. The duplex sequences chosen for these experiments (2B20, 2B39, 2B60 and 2B70, Fig. S3*A* and Table S3) were centered on a C57BL/6 mouse meiotic DSB hotspot (2B, chr9, centered at position 4447025), mapped using the SPO11-oligo approach and shown to be targeted and cut by the TOPOVIL complex *in vivo* (32). We measured TOPOVIBLΔC25 affinity for the linear DNA duplexes (from 20 to 70 bp) labeled with the ATTO647 fluorescent label using a fluorescence anisotropy-based approach. TOPOVIBLΔC25 monomers bound to 2B20 duplex DNA with a Kd of 1610 nM (Fig. 4*A*). The affinity increased (Kd = 650 nM) for the 39 bp duplex, and then progressively decreased for the 60 and 70 bp duplexes (Kd = 900 nM and 1500 nM, respectively) (Fig. 4*A*). Competition experiments performed with unlabeled 2B39 duplex confirmed that the DNA-TOPOVIBLΔC25 monomer interaction was DNA-specific and not related to the labeling with ATTO647 (Fig. S3*B*).

**Fig. 4.**
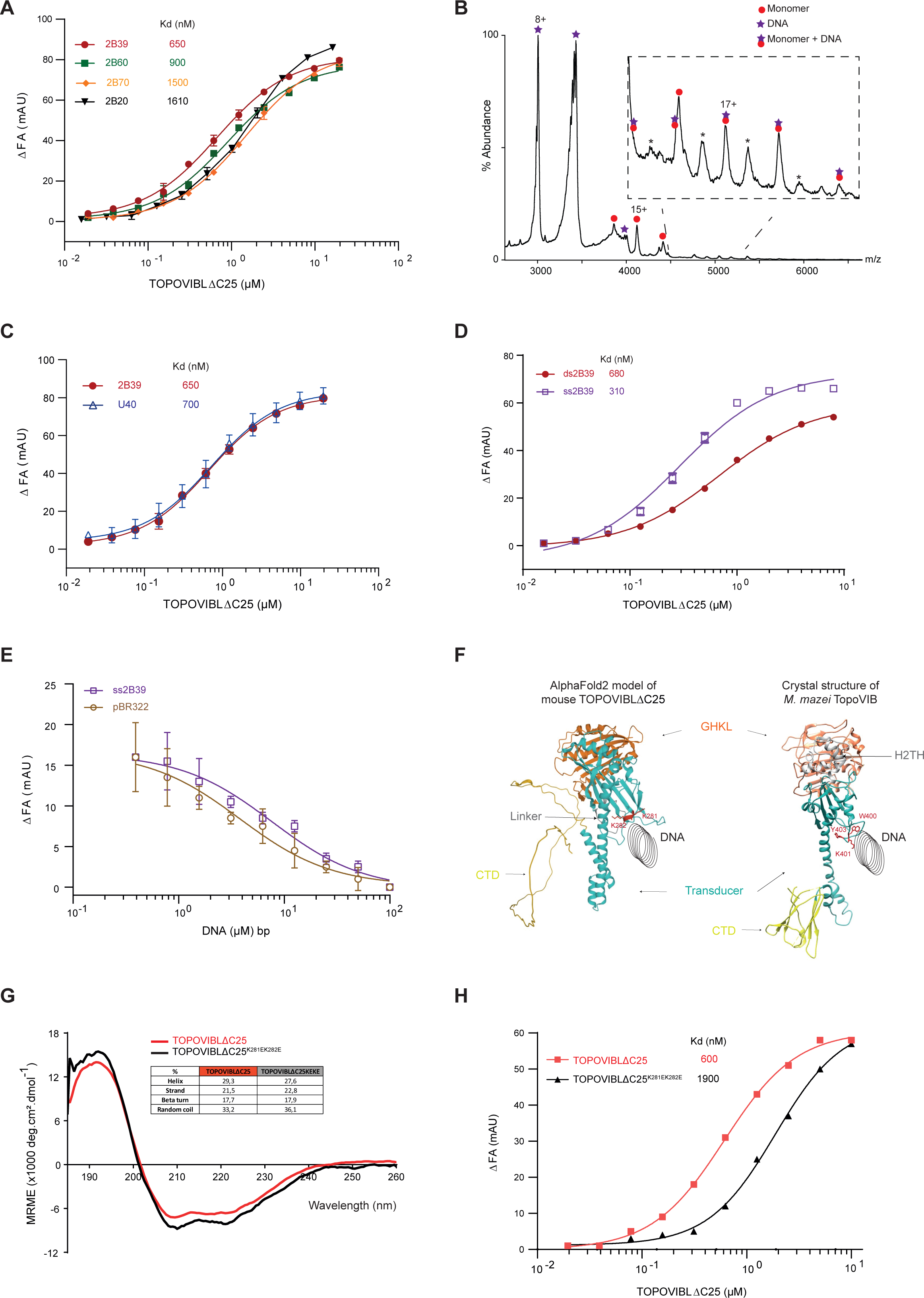
TOPOVIBLΔC25 interacts with DNA. **(*A*)** Binding of ATTO-647-labeled dsDNA fragments (related to a mouse recombination hotspot) of 20 bp (2B20), 39 bp (2B39), 60 bp (2B60) and 70 bp (2B70) to TOPOVIBLΔC25. Binding was observed as a change in fluorescence anisotropy (ΔFA) measured in milli-Anisotropy units (mAU) in function of TOPOVIBLΔC25 concentration. The points and error bars represent the mean and standard deviation of three independent experiments. **(*B*)** Native MS spectrum of an equimolar mixture of TOPOVIBLΔC25 and 2B39 duplex sequence showing the presence of the resulting 1:1 stoichiometry complex at 85,731 ± 156 Da, in addition to free monomeric TOPOVIBLΔC25 and dsDNA. **(*C)*** Binding of the ATTO-647-labeled 40 bp fragment (U40; unrelated to the mouse hotspot) to TOPOVIBLΔC25. Experiments performed as in (*A*). **(*D*)** Binding of ATTO-647-labeled 2B39 (39bp) dsDNA and ssDNA sequences to TOPOVIBLΔC25. Experiments performed as in (*A*). **(*E*)** Fluorescence anisotropy experiment assessing the ability of supercoiled DNA (pBR322) and ss2B39 to compete a fluorescein-labeled 2B39 dsDNA from TOPOVIBLΔC25. Competition was observed as a change in fluorescence anisotropy (ΔFA) as measured in milli-anisotropy units (mAU). Data are plotted as a function of the base-pair concentration (µM) of competing DNA. Points and error bars correspond to the mean and standard deviation of three independent experiments. **(*F*)** AlphaFold2 predicted structure of the mouse TOPOVIBΔC25 and crystal structure of the TopoVIB subunit of M. *mazei* TopoVI (PDB ID 2Q2E). DNA positioning is proposed. Residues mutated in the WOC motifs are highlighted in red. **(*G*)** Circular dichroism spectra of the secondary structure analysis of TOPOVIBLΔC25 (red) and TOPOVIBLΔC25^K281EK282E^ mutant protein (black). **(*H*)** Anisotropy experiment assessing the binding of an ATTO-647-labeled dsDNA (2B39) to TOPOVIBLΔC25^K281EK282E^ and TOPOVIBLΔC25. Experiment performed as in (***A***).

These experiments showed that among the different dsDNA duplexes tested, the TOPOVIBLΔC25 subunit preferentially interacted with DNA of 39 bp in length. We then used native MS to confirm the results obtained by fluorescence anisotropy and assess the stoichiometry of the TOPOVIBLΔC25 monomer-DNA interaction. The native MS spectra obtained after incubating blocked TOPOVIBLΔC25 monomers with the 2B39 duplex showed a TOPOVIBLΔC25-DNA interaction with a 1:1 stoichiometry (Fig. 4*B*). Adding DNA did not trigger the formation of TOPOVIBLΔC25 dimers in the tested conditions (Fig. 4*B*).

To investigate whether TOPOVIBLΔC25 exhibited any DNA sequence preferences, we performed similar affinity measurement using a 40 bp oligonucleotide with a sequence that did not correspond to a mouse meiotic DSB hotspot (named Unrelated 40 bp, U40). The TOPOVIBLΔC25 affinity for this duplex was similar to that for the 2B39 duplex (Fig. 4*C*). This indicated that TOPOVIBLΔC25 may not have DNA sequence preferences.

Then, we investigated whether the DNA geometry [single stranded DNA (ssDNA) *versus* double stranded DNA (dsDNA)], rather than the DNA sequence, influenced TOPOVIBLΔC25 binding to DNA, as observed for archaeal TopoVI. Affinity measurements using the fluorescence anisotropy-based approach showed that TOPOVIBLΔC25 interacted with ATTO647-labeled linear ssDNA (39 bases, ss2B39) with a Kd of 310 nM (versus 650 nM for dsDNA), indicating a better affinity for ss2B39 (Fig. 4*D*). Next, we assessed the relative affinity of TOPOVIBLΔC25 for a negative supercoiled plasmid using a competitive binding assay. We incubated TOPOVIBLΔC25 with the ATTO647-labeled 2B39 duplex and varying amounts of unlabeled ssDNA (ss2B39) or of unlabeled negatively supercoiled pBR322 plasmid. We determined the relative affinity of TOPOVIBLΔC25 for the supercoiled or ssDNA competitors by monitoring how well they interfered with the binding of linear DNA. These measurements showed that TOPOVIBLΔC25 bound to supercoiled pBR322 with a similar affinity as to 2B39 ssDNA, and thus more efficiently than to 2B39 dsDNA (Fig. 4*E*).

Altogether, we showed that the TOPOVIBLΔC25 subunit, independently of the rest of the TOPOVIL complex, directly interacts with DNA. This interaction is DNA sequence-independent, but sensitive to the DNA geometry. Specifically, TOPOVIBLΔC25 interacts preferentially with less constrained DNA (ssDNA) and with the negatively supercoiled plasmid.

### TOPOVIBLΔC25 interaction with DNA partially depends on the WOC motif

Of the three TopoVIB regions that interact with DNA (KGRR loop, WOC motif, and H2TH domain), only the WOC motif is conserved in the TOPOVIBL sequence. This motif is essential for TopoVI function and might function as an interface for robust binding to the G DNA segment (6)(7, 15). To assess whether the WOC motif function is conserved in mouse TOPOVIBL, we generated a double acidic mutation in the WOC motif of TOPOVIBLΔC25, leading to the TOPOVIBLΔC25^K281EK282E^ mutant protein (Fig. 4*F*) that was purified using the purification strategy described for TOPOVIBLΔC25. The far-UV circular dichroism and SEC results suggested that the two mutations introduced in TOPOVIBLΔC25 did not alter the protein structure (Fig. 4*G* & Fig. S3*C*). To determine whether the mutations in TOPOVIBLΔC25^K281EK282E^ affected the interaction with DNA, we performed affinity measurement by fluorescence anisotropy. Using the fluorescent-labeled 2B39 duplex, we measured a reproducible reduction of the binding affinity, with a mutant *versus* wild type Kd ratio of 3 (Fig. 4*H*). This showed that the WOC motif contributes, partly, to the contact of TOPOVIBLΔC25 with DNA. It also indicates that an additional, not conserved, part of the protein interacts with the DNA substrate.

## Discussion

Since the identification of mouse TOPOVIBL, understanding its molecular function has been a central question. Here, we provide an *in vitro* biochemical characterization of TOPOVIBL showing that it has common and also distinct features with its archaeal orthologue TopoVIB. These data help to understand its specific mode of action during meiosis.

### New structural insights on TOPOVIBL

We purified the TOPOVIBLΔC25 protein that lacks the last 25 C-terminal residues and the helix involved in the interaction with REC114 (a meiotic DSB formation accessory protein) (28). TOPOVIBLΔC25 purified as a single monomer, even if disulfide bonds can arise during the purification. The secondary structure analysis of TOPOVIBLΔC25 by circular dichroism and the structural characterization by SAXS confirmed the correct folding of the protein and experimentally validated the AlphaFold2 structure predictions. In addition, we showed that TOPOVIBLΔC25 exists in a conformational equilibrium between two states where the C-terminal disordered tail can adopt either a compact structure in close contact with the TOPOVIBLΔC25 core, or a more extended structure that is compatible with potential regulatory intramolecular interactions. This highlighted the intrinsic flexibility of the protein. The compact TOPOVIBLΔC25 arrangement could represent a closed conformation and the interaction of the missing C-terminal helix with partners (e.g. REC114) could trigger the opening of the protein by moving away the disordered part. Such conformational changes might have consequences on the TOPOVIL complex structure and activity, making it inactive (compact conformation) or active (extended conformation) for DSB formation. Adding SPO11 and REC114 for the biochemical analysis of TOPOVIBL will be essential in future studies to validate this hypothesis.

### TOPOVIBLΔC25 is not sensitive to ATP

Although the N-strap region is missing and the GHKL ATPase domain degenerated in TOPOVIBL, it is crucial to determine whether TOPOVIL activity relies on TOPOVIBL dimerization in order to understand its molecular function. Here, we experimentally confirmed that TOPOVIBLΔC25 does not interact with ATP and we did not detect its dimerization upon incubation with ATP or AMP-PNP. These results suggest that the TOPOVIBL subunit is not sensitive to ATP and that the activity of the mammalian TOPOVIL complex does not depend on TOPOVIBL subunit dimerization, although we cannot formally exclude an alternative dimerization mode. One consequence of this observation is that SPO11 dimerization and its DSB formation activity could be independent of TOPOVIBL dimerization. One possible model is that the TOPOVIBL subunit interaction with partners, such as REC114/MEI4, induces its motion and favors SPO11 dimerization and DSB formation (see below, model). Interestingly, *Arabidopsis thaliana* has two meiotic SPO11 variants (SPO11-1 and SPO11-2). Their dimerization is essential for DSB formation and depends on MTOPVIB, the TOPOVIBL orthologue (3). This observation is in accordance with the hypothesis that TOPOVIBL triggers SPO11 dimerization and activity.

### TOPOVIBLΔC25 interacts with DNA, with topological preferences

We propose that the interaction of the TOPOVIBL subunit with DNA that we reported is required for the TOPOVIL DSB meiotic activity. In Archaea, TopoVI preferentially interacts with DNA crossings and bends of supercoiled DNA. This stimulates TopoVIB dimerization and DNA cleavage. (15). TOPOVIBLΔC25 interacted with linear dsDNA, particularly of 40 bp in length. In our experimental conditions, TOPOVIBLΔC25 interacted better with smaller linear DNA than its archaeal counterpart (60 bp in Archaea) (15). The fact that DNA might interact concomitantly with the B and A subunits could explain this discrepancy. The smaller DNA size preference might reflect SPO11 missing contribution.

Moreover, TOPOVIBLΔC25 displayed better affinity for linear ssDNA and supercoiled DNA, compared to linear dsDNA. This suggests that the B subunit senses different topological statuses. As we did not detect sequence-specificity for the TOPOVIBLΔC25-DNA interaction, we might hypothesize that the DNA geometry also contributes to the substrate choice. DSB formation at meiosis onset occurs in particular accessible chromatin regions where DNA conformation may have specific features. Recent studies in yeast showed that DSB formation overlaps with specific topological regions of the genome, and in particular with regions bound by topoisomerases (33)(27, 34). This suggests a recognition of specific chromosomal features, such as negative supercoils and DNA crossings. Thus, in line with this observation, our work suggests that the TOPOVIBL subunit plays a role in sensing this specific topological status. The DNA topological status might also influence TOPOVIL activity by triggering TOPOVIBL motion that could favor the formation of an active TOPOVIBL-SPO11 hetero-tetramer *in vivo*.

In Archaea, three TopoVIB domains interact with DNA (15), but only the WOC motif (or Stalk/WKxY) is conserved in the mouse protein. We found a partial contribution of the TOPOVIBL WOC motif to the TOPOVIBLΔC25-DNA interaction. Different non-exclusive hypotheses might account for this difference. First, the double mutation introduced in the WOC domain of TOPOVIBLΔC25 (two lysine residues replaced by glutamic acid residues) had no detectable impact on the protein structure (Fig. 4*F*-*G* & Fig. S3*C*). In Archaea, the generated mutants targeted a tryptophan, a lysine and a tyrosine residue within the WOC motif that presumably affect more strongly the protein structure and the DNA binding affinity (15). This might partly explain why the archaeal mutant has a clearer effect. Second, the mild contribution of the WOC motif in the interaction with DNA indicates that additional parts of the protein, not conserved in Archaea, are implicated in the interaction with DNA, suggesting an evolution of the regulatory role of the TOPOVIBL-DNA interaction in the regulation of the meiotic enzyme activity. Interestingly, in yeast, an interaction between the Rec102 subunit of the DSB core complex (equivalent to the transducer region of TOPOVIBL) and DNA was reported (23). However, the Rec102 interacting region is not conserved and its biological significance remains to be studied.

Altogether, our results, in line with previous studies, highlight the importance of the TOPOVIBL-DNA interaction. They also show that this interaction depends on a specific DNA architecture and is partly mediated by a conserved motif and possibly also by additional (not identified yet) regions of the protein or of the complex.

### A proposed mode of action for the TOPOVIL complex

Based on our results and previous studies, we propose the following model for TOPOVIL molecular activity (Fig. 5). First, the SPO11-TOPOVIBL dimeric complex interacts with specific DNA structures (e.g. negative supercoiled or DNA crossing) (Step 1, Fig. 5). Then, the SPO11 subunit dimerizes. Different non-exclusive parameters might promote and stabilize this dimerization. The DNA topological environment sensed by TOPOVIBL could be one of these parameters. The MEI4/REC114 (1:2) complex also might act as a clamp to hold together the two SPO11 subunits (24–26) (28). This clamping activity could be mediated by the C-terminal helix of TOPOVIBL that interacts with REC114 and possibly by TOPOVIBL dynamic conformation revealed by our SAXS data (Step 2, Fig.5). In this model, the open and compact conformations of TOPOVIBL allowing interaction or not with REC114, might represent an active and inactive state for TOPOVIL, respectively. Accumulation of proteins that create a matrix and a condensate-like environment to favor protein-protein interactions also might favor SPO11 dimerization, as proposed in alternative studies (24, 26, 27)(27). As observed in archaea, the SPO11 dimerization could then induce conformational changes of this catalytic subunit, making it efficient for DSB formation activity (Step 3, Fig. 5). Finally, following cleavage, SPO11 binding interfaces are destabilized. These structural changes could be associated to a compact conformation of TOPOVIBL and to the destabilization of MEI4/REC114 interaction. Altogether, this leads to the release of the two ends of the DSB (Step 4, Fig. 5).

**Fig. 5.**
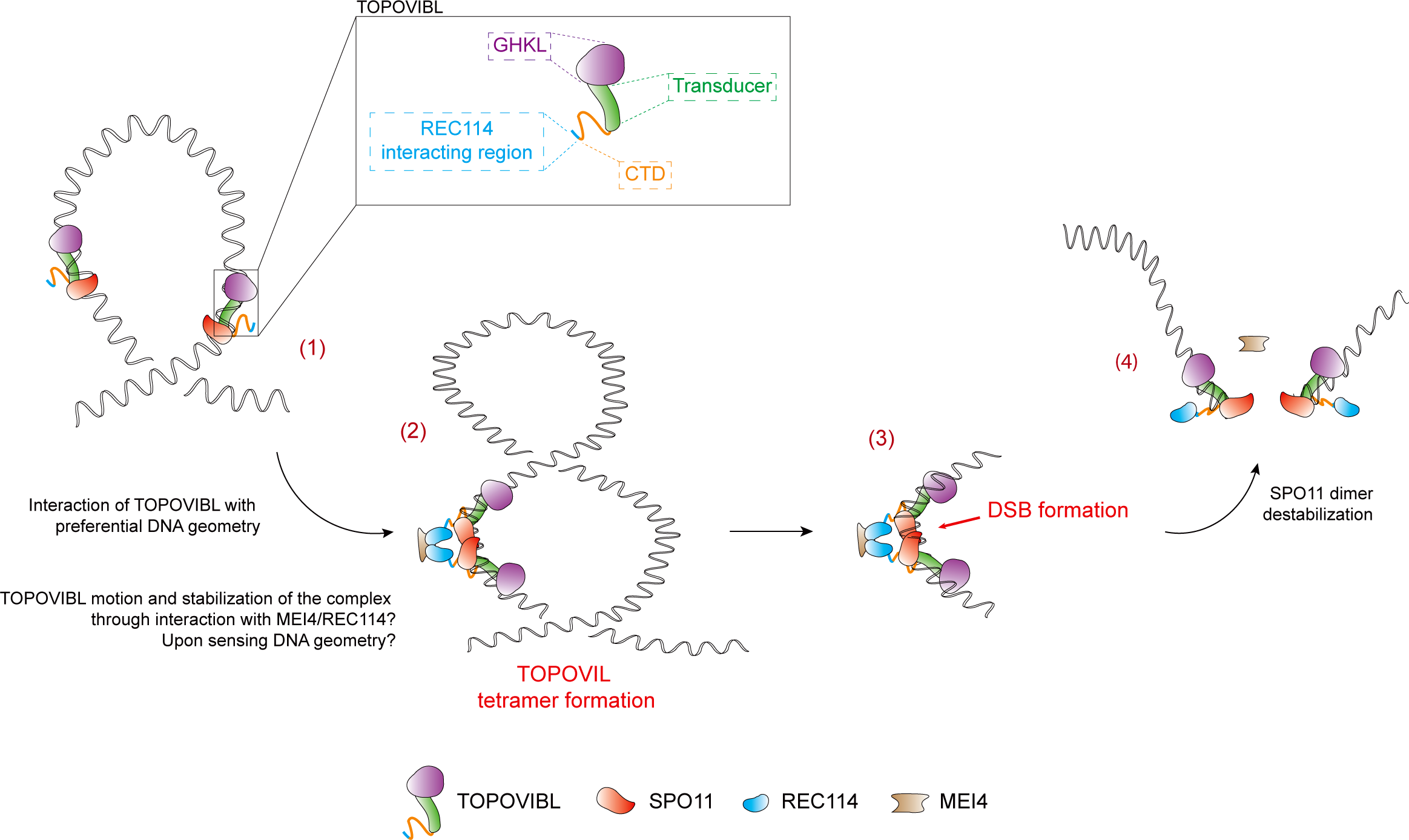
Model for TOPOVIL mode of action. (1) The TOPOVIBL-SPO11 complex interacts with DNA, preferentially with supercoiled or ssDNA. (2) TOPOVIBL-SPO11 complex dimerizes through SPO11-SPO11 interactions, forming a hetero-tetramer. DNA topology or interaction with additional partners: REC114/MEI4 (2:1) complex, can mediate the hetero-tetramer formation. (3) SPO11 dimerization is compatible with the formation of DSB. (4) DSB formation leads to the destabilization of the SPO11 interface, and to TOPOVIBL conformational changes that release the REC114/MEI4 complex and to the formation of irreversible DNA DSBs.

This study offers a first step in the *in vitro* characterization of TOPOVIBL molecular function and helps to understand how it regulates meiotic DSB formation activity in mammals. This study also paves the way to additional *in vitro* characterization studies particularly of the full TOPOVIL complex (TOPOVIBL and SPO11), with accessory factors (MEI4/REC114), to fully characterize how it is regulated by TOPOVIBL.

## Materials and methods

### Plasmid constructs for protein production

The MBP-TOPOVIBLΔC25 protein was produced in Sf9 insect cells using the bac-to-bac baculovirus expression system (Invitrogen). Mouse *Top6bl* cDNA was first amplified from the pl159 plasmid (Table S4) (pETDuet-His-Flag-GM960) using the Oli246 and Oli237 primers (Table S3) (6). After replacing the *Nhe*I restriction site by the *Kpn*I site in the pl207 plasmid (pFastBac-MBP-Sae2-10His)(35), the amplified *Top6bl* gene was cloned in the pl207 plasmid, at the place of the *Sae2* gene, using the *Kpn*I and *Xma*I restriction sites to generate the pl237 plasmid (pFastBac-MBP-TOPOVIBL). pl237 was used as a template for mutagenic PCR using Oli329 and Oli330 primers to produce the pl397 plasmid (pFastBac-MBP-TOPOVIBLΔC25). Bacmid 353, used for the production of MBP-TOPOVIBLΔC25 in insect cells, was subsequently produced from pl397 according to the manufacturer’s instructions (Bac-to-Bac Baculovirus Expression system, Invitrogen). The pl485 plasmid (pFastBac-MBP-TOPOVIBLΔC25^K281EK282E^) was generated by the GeneScript company using pl397 as template. Bacmid 485, used for the production of MBP-TOPOVIBLΔC25^K281EK282E^ in insect cells, was generated according to the manufacturer’s instructions (Bac-to-Bac Baculovirus Expression system, Invitrogen) using pl485.

### TOPOVIBLΔC25 and TOPOVIBLΔC25K281EK282E production and purification

Sf9 cells (4.10^6^/ml) were infected with the P3 baculovirus at a 1:200 ratio and grown in EX-CELL 420 serum-free medium (14420C, Merk). After 48 hours of incubation at 28°C, 180 rpm rotation, cells were harvested and resuspended in lysis buffer (20 mM HEPES pH 7.5, 500 mM KCl), supplemented with 0.5x protease inhibitor cocktail (5056489001, Roche) (45 ml lysis buffer for 1 liter culture). Lysis (25 ml cell suspension) was performed on a Vibra-Cell 72405 sonicator (Bioblock Scientific) with a small probe (630-0422, Thermo Fisher Scientific), using 50% amplitude, 2.5 seconds pulse for 3 minutes. Whole cell extracts were centrifuged and supernatants were filtered through 0.45 µm filters (146561, Clearline). Proteins were subsequently purified on amylose resin (E8021L, NEB): 3 ml resin per column and 45 ml of extract. The resin was washed and resuspended in 3 ml of lysis buffer, supplemented with a homemade GST-tagged 3C protease (PreScission) (0.033 mg/ml final concentration) to remove the MBP tag by overnight incubation at 4°C. The cleaved fractions were collected by gravity flow as the flow through of the column and beads were resuspended in 1.5 ml of lysis buffer with 3C protease for a second round of cleavage. After 2 hours on ice, the second cleaved fraction was collected. The resin column was then eluted with lysis buffer containing 10 mM maltose, for evaluation of the non-cleaved fraction that remains in the column. The cleaved fractions were pooled and incubated with glutathione Superflow agarose (25237, Thermo Fischer Scientific) to remove the GST-tagged 3C protease. The flowthrough was collected as the 3C protease-depleted fraction, concentrated on an Amicon ultra-15 centrifugal filter unit 30.000 MWCO (UFC903024, Millipore), and injected onto a Superdex 200 10/300 column (GE Healthcare) for Size-Exclusion Chromatography (SEC) on a High-Performance Liquid Chromatography (HPLC) ÄKTA Pure system (GE Healthcare). The column was equilibrated and eluted with the lysis buffer. For protein purification in the presence of 2-iodoacetamide (IAA), lysis buffer was supplemented with 10 mM IAA (I6125, Sigma Aldrich).

### Tandem mass spectrometry (MS/MS) of purified TOPOVIBLΔC25

Samples were digested essentially according to the protocol described by Shevchenko (36). Samples were loaded on 10% SDS-PAGE gels for short migration (Mini-Protean TGX Precast gels, Bio-Rad). One band was cut after stacking migration. Proteins were in-gel digested using 1µg trypsin (Trypsin Gold, Promega)(37). The obtained peptides were analyzed online using a Q-Exactive HF-X mass spectrometer (Thermo Fisher Scientific) interfaced with a nano-flow HPLC (RSLC U3000, Thermo Fisher Scientific). Samples were loaded onto a 25 cm reverser phase column (Acclaim Pepmap 100^®^, NanoViper, Thermo Fisher Scientific) and separated using a 60-min gradient of 6 to 40% of buffer B (80% acetonitrile, 0.1% formic acid) at a flow rate of 300 nL/min. MS/MS data were analyzed in data-dependent mode (Xcalibur software 4.1, Thermo Fisher Scientific). Full scans (350-1500 m/z) were acquired in the Orbitrap mass analyzer at a resolution of 120,000 m/z resolution at 200 m/z. The twenty most intense ions (charge states ≥2) were sequentially isolated and fragmented by high-energy collisional dissociation in the collision cell and detected with a resolution of 15,000 resolution. Spectral data were analyzed using the MaxQuant software with default settings (v1.6.10.43)(38). All MS/MS spectra were analyzed with the Andromeda search engine against a decoy database that included the TOPOVIBLΔC25 protein sequence, AcNMPV proteome (UP000008292), *Spodoptera frugiperda* proteome (Taxonomy 7108) (downloaded from UniProt, release 2021_01, https://www.uniprot.org/) and classical contaminants. Default search parameters were used, with Oxidation (Met) and Acetylation (N-term) as variable modifications and Carbamidomethyl (Cys) as fixed modification. The false discovery rate (FDR) was set to 1% for peptides and proteins. A representative protein ID in each protein group was automatically selected using an in-house bioinformatics tool (Leading_v3.4). First, proteins with the most numerous identified peptides were isolated in a “match group” (proteins from the “Protein IDs” column with the maximum number of “peptides counts”). For the match groups where more than one protein ID were present after filtering, the best annotated protein in UniProtKB, release 2021_01 (reviewed rather than automatic entries, highest evidence for protein existence) was defined as the “leading” protein. The iBAQ value was used to highlight the most abundant protein in the sample.

### Circular Dichroism

The protein secondary structure was determined by far UV-circular dichroism spectroscopy using a Chirascan instrument (Applied Photophysics). Protein samples (0.2 - 0.6 mg/ml in 20 mM HEPES pH 7.5, 50 mM KCl) were loaded in quartz cuvettes, 0.1 mm path length (Hellma), and the wavelength scans were obtained between 185 and 260 nm, step size 0.5 nm, bandwidth 1 nm. Three different measurements were done for each sample. The mean value of the three measurements was buffer-corrected and converted to mean molar residue ellipticity [θ] (deg×cm2/dmol). Data deconvolution, to determine the secondary structure contents, was done with the CDNN software.

### Size exclusion chromatography coupled to multi-angle light scattering (SEC-MALS)

Gel filtration-purified protein samples (1 - 5 mg/ml) were analyzed by SEC-MALS. Protein samples (20µl) were injected onto a Superdex 200 10/300 Increase column (GE Healthcare) equilibrated with 20 mM HEPES pH 7.5, 500 mM KCl. SEC was performed on a HPLC system (Agilent Infinity II). Multi-angle light scattering was detected with a miniDAWN TREOS instrument (Wyatt Technology) and refractometry measured with an Optilab T-Rex refractive index detector (Wyatt Technology). SEC-MALS data were collected and analyzed with the ASTRA 7 software (Wyatt Technology).

### Native mass spectrometry

Before MS analysis, proteins were buffer-exchanged against 100 mM ammonium acetate buffer pH 7.4 (Sigma) using Amicon centrifugal filters (Merck) followed by Bio-Spin microcentrifuge columns (Bio-Rad Laboratories). 2B39 DNA was buffer-exchanged using Bio-Spin microcentrifuge columns (Bio-Rad Laboratories). Intact MS spectra were recorded on a Synapt G2-Si HDMS instrument (Waters Corporation) modified for high mass analysis and operated in ToF mode. Samples were introduced into the ion source using borosilicate emitters (Thermo Scientific). Optimized instrument parameters were as follows: capillary voltage 1.4 kV, sampling cone voltage 100 V, offset voltage 120 V, transfer collision voltage 15 V, argon flow rate 8 mL/min and trap bias 5 V. Trap collision voltage varied between 50 and 120 V. Data were processed with MassLynx v.4.2 (Waters).

### TOPOVIBLΔC25 dimer incubation with DTT

SEC-purified TOPOVIBLΔC25 dimers (200 µg in 40 µl 20 mM HEPES pH 7.5, 500 mM KCl) were incubated on ice with 50 mM DTT (Euromedex, EU0006), final concentration, for 30 min. Then, samples were injected on a Superdex 200 10/300 Increase column (GE Healthcare) for SEC on a HPLC ÄKTA Pure system (GE Healthcare) in lysis buffer containing 1 mM DTT.

### TOPOVIBLΔC25 dimer incubation with TCEP

12 µg of SEC-purified TOPOVIBLΔC25 monomers or dimers, in 30 µl of 20 mM HEPES pH 7.5, 500 mM KCl, were incubated on ice with different TCEP concentrations for 30 min. Then, samples were analyzed by non-reducing-PAGE and Coomassie staining.

### Small-angle x-ray scattering (SAXS) and ensemble optimization of TOPOVIBLΔC25

SAXS measurements were performed at the SWING beamline, SOLEIL synchrotron (Saint-Aubin, France), using a wavelength of 1.0332 Å and sample-to-detector distance of 2 m. Before data collection, the TOPOVIBLΔC25 dimer sample was incubated on ice with 2 mM of DTT (final concentration) for 1 hour. After incubation, a volume of 50 µL from the 12 mg/mL sample was injected onto a Superdex 200 Increase 10/300 GL column, running at a flow rate of 0.5 mL/min. The column was pre-equilibrated with sample buffer (20 mM HEPES pH 7.5, 500 mM KCl, and 5% glycerol supplemented with 1 mM DTT). SAXS 2D images were azimuthally averaged into a 1D scattering intensity curve using the in-house Foxtrot software (available at https://www.synchrotron-soleil.fr/en/beamlines/swing). The scattering frames from buffer and sample were selected and averaged using CHROMIXS, after a buffer subtraction step. The obtained SAXS intensity curves were analyzed using PRIMUS from the ATSAS software package (39). The collected scattering vector ranges ranged from q=0.011 to 0.055 Å–1 (q=4π/λ sin θ where 2θ is the scattering angle and λ the X-ray wavelength) (40). To represent the flexibility present in TOPOVIBLΔC25 monomers, an ensemble of 10,000 conformations was modeled using RANCH from the Ensemble Optimization Method (EOM)(30). For that, the AlphaFold2 model (af-j3qmy9-f1) lacking the C-terminal region (C451 to S579) was used as input reference, and was then modeled as a random-coil (from C451 to P554). The theoretical scattering intensities were calculated for all conformations. Then, sub-ensembles of 50 conformations were selected to collectively fit the experimental SAXS data.

### TOPOVIBLΔC25 dimerization test in the presence of ATP or AMP-PNP

20 µM of TOPOVIBLΔC25 monomers purified in the presence of IAA was incubated with 50 µM ATP (P0756L, NEB) or AMP-PNP (64473120, Roche Diagnostics) on ice for 30 min. Then, samples underwent SEC on a Superdex 200 10/300 Increase column (GE Healthcare) on a HPLC ÄKTA Pure system (GE Healthcare). The SEC fractions were analyzed by SDS-PAGE.

### Conversion of TOPOVIBLΔC25 dimers to monomers using DTT and IAA

As the dimeric form displayed higher purity than the monomeric form, the dimeric form was converted into a blocked monomeric form by incubation with DTT and IAA. SEC-purified TOPOVIBLΔC25 dimers (1.5 - 2 mg) were incubated with 50 mM DTT on ice for 30 min. Then, iodoacetamide (IAA), 10 mM final, was added to the sample, and incubated on ice for 30 min. The sample was centrifuged at 14.000 g, 4°C, for 10 min, and injected onto a Superdex 200 10/300 Increase column (GE Healthcare) on a HPLC ÄKTA Pure system (GE Healthcare) for SEC. The fractions of the peak center that corresponded to TOPOVIBLΔC25 monomers were used for native MS and the dimerization test in the presence of ATP or AMP-PNP.

### Fluorescence anisotropy and competition assay

The DNA sequences used for fluorescence anisotropy corresponded to a mouse C57BL/6 meiotic double-strand break (DSB) hotspot (2B, chr9, centered on the nucleotide at position 4447025, GRCm38/3310)(32). The 40 bp unrelated sequence (U40) was chosen from the literature: 40 bp duplex from (15). The ATTO647-labeled single-stranded DNA fragments and the complementary strands were ordered from Integrated DNA Technologies. Double-stranded DNA (2B20, 2B39, 2B60 and 2B70) were produced by mixing 4µl of 100 µM labeled DNA with 4.4 µL of the complementary strands at 100 µM (to ensure that all labeled DNA was hybridized) in 50 mM NaCl, 20 mM HEPES pH 7.5, heated at 95 °C for 20 min, and cooled down at room temperature for 4 hours. Hybridization was checked on a 4-12% Bis-Tris gel in 1X TAE.

Fluorescence anisotropy experiments were performed in the following conditions. Purified TOPOVIBLΔC25 was dialyzed against binding buffer (50 mM HEPES pH 7.5, 250 mM potassium glutamate, 5% glycerol) to reduce the salt concentration compared with the protein purification buffer. Then, samples were serially diluted in two-fold steps in binding buffer and incubated with the ATTO647-labeled DNA substrate, either single strand (ss) or double strand (ds), to obtain a final concentration of 20 nM, in the dark and on ice for 5 min. Reactions were diluted to the final binding assay conditions (10 mM HEPES pH 7.5, 50 mM potassium glutamate, 5% glycerol), and incubated on ice for 10 min. Fluorescence anisotropy was measured at ambient temperature using a Safire microplate reader (TECAN) with excitation at 635 nm and emission at 680 nm. Data were the mean of at least three independent experiments, and were fitted using a sigmoidal dose-response model (GraphPad Prism, GraphPad software).

Competition assays were carried out as described for the direct binding assay, with protein samples diluted in binding buffer to a final concentration of 1.3 µM and incubated with the ATTO647-labeled 2B39 dsDNA at a final concentration of 20 nM. Unlabeled 2B39 ssDNA or the negatively supercoiled pBR322 plasmid (ThermoScientific) were titrated by serially diluting them in two-fold steps in the final binding assay buffer, starting from 100 µM bp. Incubation conditions were the same as for the direct titration. Anisotropy data were fitted with an on-site competition equation.

## Acknowledgement

The Centre for Structural Biology (CBS) is a member of France-BioImaging (FBI) and the French Infrastructure for Integrated Structural Biology, two national infrastructures supported by the French National Research Agency (grant nos. ANR-10-INBS-04-01 and ANR-10-INBS-05, respectively). Mass spectrometry experiments were carried out using the facilities of the Montpellier Proteomics Platform (PPM, BioCampus, Montpellier) that is supported by FEDER/Région Occitanie, MUSE and the Labex EpiGenMed. We thank our colleagues for helpful discussions: Pau Bernado for SAXS data analysis, Raphael Guerois for bioinformatic analysis, William Bourguet and Frederic Baudat for discussion on the manuscript and all lab members. This work was supported by CNRS INSERM ATIP-Avenir 2017 program and ANR CONDENSin3R (ANR-20-CE12-0016-02) grants. I.T. is supported by a grant from the ANR (ANR-21-CE11-005-01). We thank the SWING beamline at the SOLEIL synchrotron, Saint-Aubin, France, for beamtime allocation to the project and assistance during data collection. BdM was funded by CNRS, European Research Council (ERC) Executive Agency under the European Union Horizon 2020 research and innovation programme (Grant Agreement no. 883605).

## Figure legends

**Fig. S1.**
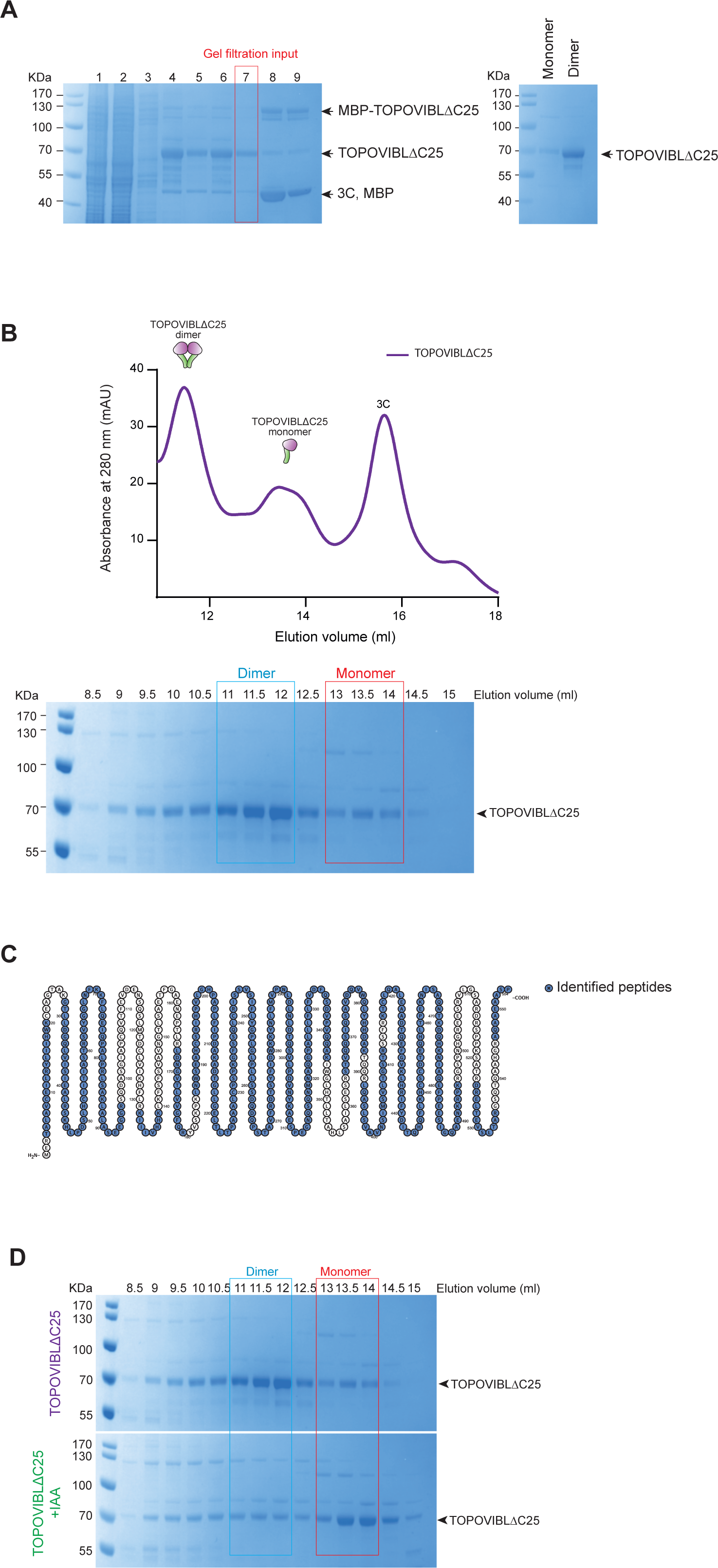
**(*A*)** SDS-PAGE and Coomassie staining of the samples obtained from the different TOPOVIBLΔC25 purification steps. Left panel: gel of the samples obtained from cell lysis to the SEC input preparation. Lane 1: Sf9 cell lysate, 2: MBP column flow through (FT), 3: MBP column wash, 4: Cleaved fraction after the first incubation with the PreScission (3C) enzyme, 5: Cleaved fraction after the second PreScission incubation, 6: FT of the cleaved fractions after incubation with glutathione beads, 7: Gel filtration input obtained from fraction 6 after concentration on Amicon 30.000 MWCO, 8: Elution 1 from the amylose column, 9: Elution 2 from the amylose column. Right panel: gel of the SEC pooled fractions corresponding to the peaks of the monomer and the dimer forms after concentration. **(*B*)** SEC chromatogram profile (upper) and SDS-PAGE analysis (lower) of fractions after TOPOVIBLΔC25 affinity purification and tag removal on the amylose column. Upper panel: SEC elution profile of a Superdex 200 10/300 column. The first peak, at 11-12 ml elution volume, is compatible with TOPOVIBLΔC25 dimers, the second peak, at 13-14 ml elution volume, with monomers, and the third peak, at 15.7 ml elution volume, corresponds to the PreScission protease (3C). Lower panel: SDS-PAGE analysis of SEC elution fractions followed by Coomassie staining. The fractions corresponding to the dimer peak are delimited by a light blue box and the fractions of the monomer peak by a red box. **(*C*)** Coverage of the peptides identified by MS/MS analysis of the SEC-purified TOPOVIBLΔC25 protein. The peptides identified in the MS/MS analysis are in blue. **(*D*)** SDS-PAGE and Coomassie staining of the SEC elution fractions of TOPOVIBLΔC25 purified with (bottom gel) or without IAA (top gel) in the cell lysis buffer. The fractions corresponding to the dimer peak are delimited by a light blue line and the fractions of the monomer peak by a red line.

**Fig. S2:**
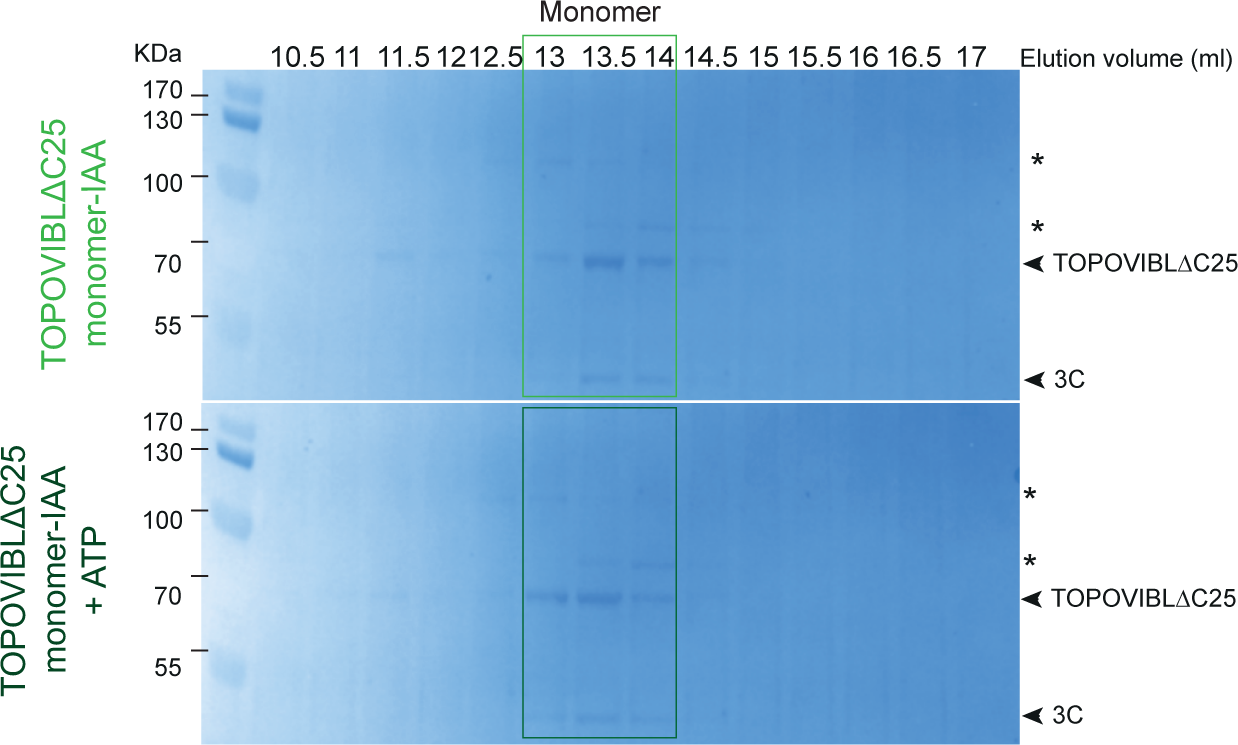
SDS-PAGE and Coomassie staining of the SEC elution fractions of TOPOVIBLΔC25 monomers (top gel) and TOPOVIBLΔC25 monomers incubated with ATP (bottom gel). In both cases, TOPOVIBLΔC25 elutes mainly in monomeric form (fractions 13 to 14 ml; light green box and dark green box in the top and bottom gels respectively). IN: SEC input. *: contaminant bands.

**Fig. S3:**
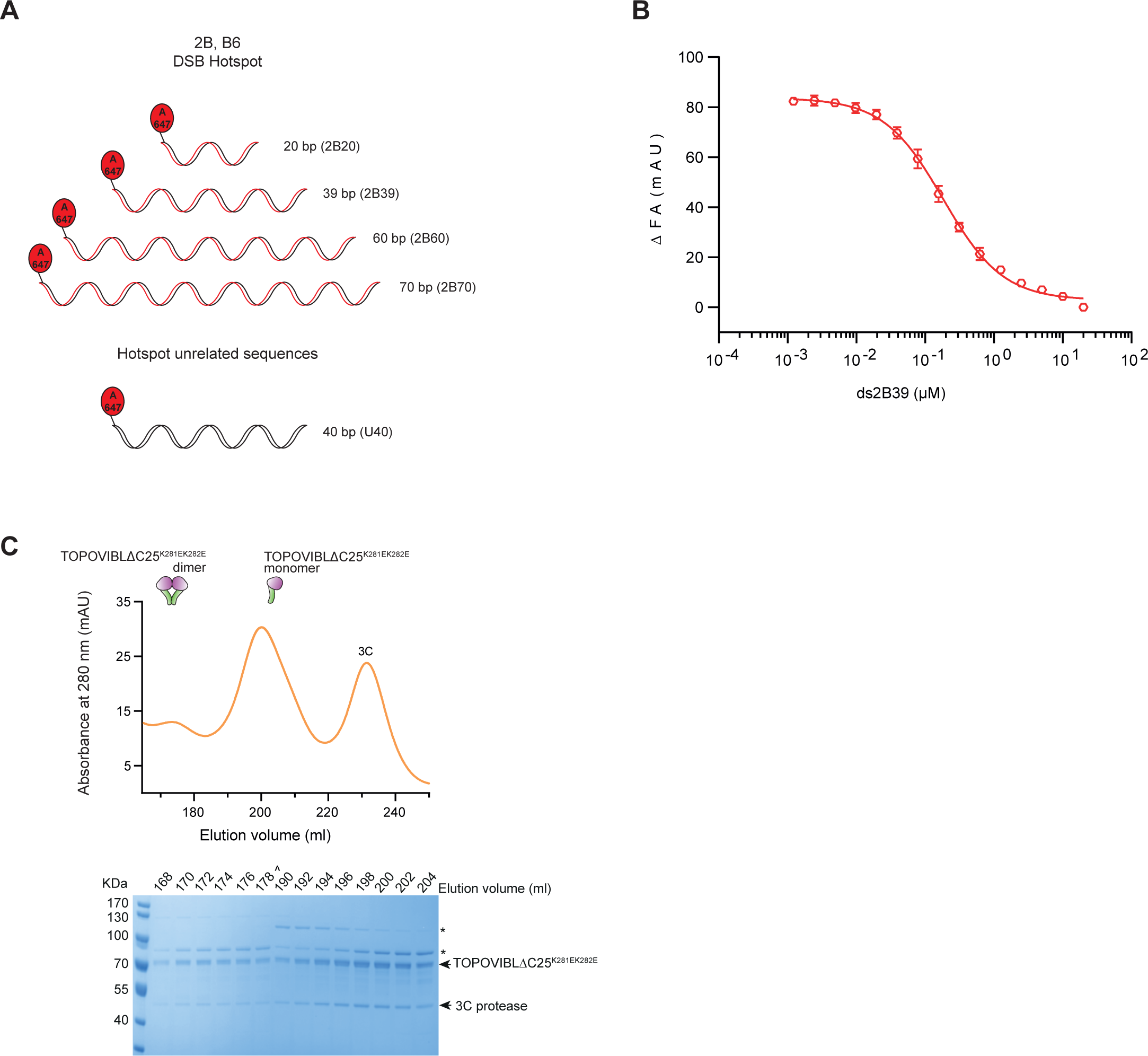
**(A)** Schematic representation of the DSB hotspot-related (2B20, 2B39, 2B60, 2B70) and -unrelated (U40) dsDNA. **(B)** Anisotropy experiment to measure the ability of unlabeled 2B39 ssDNA to compete with ATTO-647-labeled 2B39 dsDNA (20 nM) for binding with TOPOVIBLΔC25 (1.3 µM). Binding was observed as a change in fluorescence anisotropy (ΔFA) measured in milli-Anisotropy units (mAU) in function of the unlabeled DNA concentration. **(C)** SEC chromatogram and SDS-PAGE analysis of fractions after TOPOVIBLΔC25^K281EK282E^ affinity purification and tag removal on an amylose column. Upper panel: SEC elution profile using a Superdex 200 26/600 column. The main elution peak at 200 ml corresponds to TOPOVIBLΔC25^K281EK282E^ monomers. Lower panel: SDS-PAGE analysis and Coomassie staining of the SEC elution fractions. Arrows indicate TOPOVIBLΔC25^K281EK282E^ and the PreScission protease (3C). *: contaminants.

**Supplementary Table 1:** The TOP 10 highest MS/MS count proteins identified in the MS/MS analysis of the SEC-purified TOPOVIBLΔC25 sample.

| Protein IDs | DESC_CHECK | Peptides | Mol. weight [kDa] | Intensity | iBAQ | MS/MS count |
| --- | --- | --- | --- | --- | --- | --- |
| P001_TOP6deltaC25 | TOPOVIBLΔC25 | 40 | 61,165 | 51257000000 | 1708600000 | 709 |
| A0A2H1WFA2 | Heat shock 70 kDa protein cognate 4 (Fragment) | 45 | 71,507 | 2983300000 | 93227000 | 118 |
| FPPCON_P00761 | Contaminant: Trypsin | 8 | 24,409 | 17286000000 | 1571400000 | 84 |
| A0A2H1WFD3;A0A2H1WFD2 | Tubulin beta chain | 14 | 49,567 | 351850000 | 17593000 | 38 |
| A0A2H1WYZ0;G3CKA7;A0A2H1V6C6;A0A2H1WFF5;A0A2H1WZY3 | Tubulin alpha chain | 12 | 49,936 | 437440000 | 20831000 | 31 |
| P41473 | Uncharacterized 30.6 kDa protein in IAP2-VLF1 intergenic region | 15 | 30,567 | 552390000 | 34524000 | 30 |
| A0A2H1WX17;Q8I866 | SFRICE016243.2 (Fragment) | 21 | 73,122 | 435130000 | 14504000 | 28 |
| Q8WQJ2 | 60S acidic ribosomal protein P0 | 10 | 33,957 | 191620000 | 13687000 | 21 |
| A0A2H1V5C0 | T-complex protein 1 subunit delta | 11 | 56,888 | 326750000 | 10211000 | 20 |
| A0A2H1W0E4;V5ND41 | SFRICE014504.2 | 12 | 45,433 | 225470000 | 11273000 | 18 |

**Supplementary Table 2:** Protein species from native MS analysis.

| Protein species | Theoretical MW (Da) | Experimental MW (Da) |
| --- | --- | --- |
| TOPOVIBLΔC25 |  |  |
| Dimer | 122 639 | 123 044 ± 24 |
| Open Dimer |  | 122 908 ± 76 |
| Blocked Monomer | 61946 | 61 844 ± 55 |
| Open Blocked Monomer |  | 61 710 ± 8 |
| Contaminants |  |  |
| Hsp70 Monomer | 69,875 / 73108.83 | 71 448 ± 7 |
| Hsp70 Dimer |  | 142 875 ± 7 |

**Supplementary Table 3:**
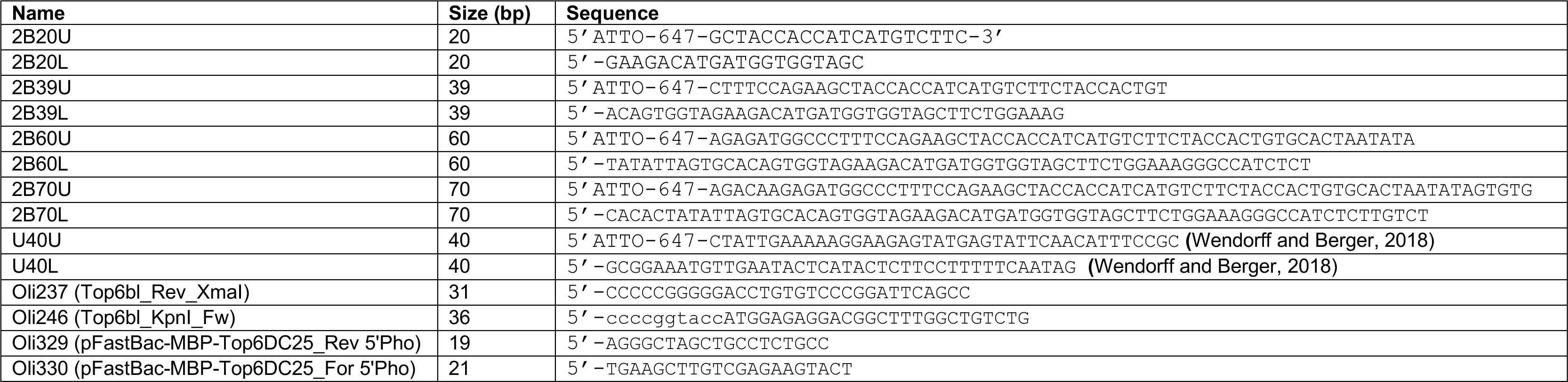
Oligonucleotides.

**Supplementary Table 4:** Plamsids.

| Name | Description |
| --- | --- |
| pI159 | pETDuet-His-Flag-GM960 |
| pI207 | pFastBac_MBP_Sae2_10His |
| pI237 | pFastBac-MBP-TopoVIBL |
| pI397 | pFastBac-MBP-TOPOVIBLΔC25 |
| pI485 | pFastBac-MBP-TopoVIBLΔC25<sup>K281EK282E</sup> |

